# Construction of Fosmid-based SARS-CoV-2 replicons for antiviral drug screening and replication analyses in biosafety level 2 facilities

**DOI:** 10.1101/2023.02.15.528742

**Authors:** Shunta Takazawa, Tomohiro Kotaki, Satsuki Nakamura, Chie Utsubo, Masanori Kameoka

## Abstract

The coronavirus disease 2019 (COVID-19) pandemic, caused by severe acute respiratory syndrome coronavirus-2 (SARS-CoV-2), has necessitated the global development of countermeasures since its outbreak. However, current therapeutics and vaccines to stop the pandemic are insufficient and this is mainly because of the emergence of resistant variants, which requires the urgent development of new countermeasures, such as antiviral drugs. Replicons, self-replicating RNAs that do not produce virions, are a promising system for this purpose because they safely recreate viral replication, enabling antiviral screening in biosafety level (BSL)-2 facilities. We herein constructed three pCC2Fos-based RNA replicons lacking some open reading frames (ORF) of SARS-CoV-2: the Δorf2-8, Δorf2.4, and Δorf2 replicons, and validated their replication in Huh-7 cells. The functionalities of the Δorf2-8 and Δorf2.4 replicons for antiviral drug screening were also confirmed. We conducted puromycin selection following the construction of the Δorf2.4-puro replicon by inserting a puromycin-resistant gene into the Δorf2.4 replicon. We observed the more sustained replication of the Δorf2.4-puro replicon by puromycin pressure. The present results will contribute to the establishment of a safe and useful replicon system for analyzing SARS-CoV-2 replication mechanisms as well as the development of novel antiviral drugs in BSL-2 facilities.

## 1. Introduction

Severe acute respiratory syndrome coronavirus-2 (SARS-CoV-2) is an etiological pathogen of coronavirus disease 2019 (COVID-19) (Kung et al., 2022). Since its outbreak in 2019, this infectious disease has caused a worldwide pandemic. Patients with COVID-19 commonly have fever, cough, fatigue, and shortness of breath, and severe cases may develop acute renal failure, acute respiratory distress syndrome, septic shock, and, ultimately, death (Chen et al., 2020; Huang et al., 2020). As of early January 2023, more than 660 million individuals have been infected and more than 6.7 million have died, which has had a huge impact on not only the health care system, but also the world economy (CSSE-JHU, 2023).

Extensive efforts have been made to combat SARS-CoV-2. Novel therapeutics, including monoclonal antibodies (mAb) and small compounds, and vaccines using new platforms, such as mRNA and viral vector vaccines, have been developed (Atluri et al., 2022; CDC, 2022). However, their efficacy is limited. Few therapeutic options are currently available for severe cases; anti-inflammatory agents are mainly used and the antiviral drug, remdesivir has limited efficacy (Atluri et al., 2022). Oral drugs, such as molnupiravir and paxlovid, have also been developed for outpatients, but are not very helpful because of low efficacy (molnupiravir) and adverse drug reactions (paxlovid) (Atluri et al., 2022). Moreover, the effects of therapeutic mAb may be evaded by amino acid mutations in the spike protein of SARS-CoV-2. The efficacy of some mAb therapeutics against the Omicron variant was markedly reduced due to the presence of more than ten amino acid alterations in its spike gene (Takashita et al., 2022). These mutations have also affected the effectiveness of vaccines, and breakthrough infections, which occur after vaccination, are frequently reported (Kuhlmann et al., 2022). In addition to spike mutations, other mutations in viral proteins that are the target of antiviral agents, such as SARS-CoV-2 3C-Like protease (3CLpro), have been suggested to affect the clinical outcomes of drug therapeutics (Heilmann et al., 2023). In this context, countermeasures against COVID-19 are insufficient, which necessitates the further development of vaccines and antivirals.

The development of an effective and safe system to perform research on SARS-CoV-2 is urgently needed because SARS-CoV-2 is a transmissible pathogen that requires a biosafety level (BSL)-3 facility for its handling. Replicons, which are self-replicating RNAs, are a promising system to overcome this obstacle. Replicons possess genes that are involved in replication, but lack those required to produce infectious virions. Since replicons only mimic viral replication, they enable the search for compounds that exert inhibitory effects on SARS-CoV-2 replication and analyses of viral replication mechanisms, even in a BSL-2 facility.

SARS-CoV-2 belongs to the genus betacoronavirus and its genome possess approximately 30 kbp positive-sense single-stranded RNA (Kung et al., 2022). The genome comprises 12 open reading frames (ORF), and ORF1, which codes non-structural proteins that are essential for viral replication, is a pivotal component of replicons (Kung et al., 2022). We previously constructed a replicon (named here as the Δorf2-8 replicon) that lacked the genes encoding ORF2 [spike (S)], ORF3, ORF4 [envelope (E)], ORF5 [membrane (M)], ORF6, ORF7, and ORF8, and possessed only 3 ORFs: ORF1, ORF9 [nucleocapsid (N)] with HiBiT fused at the C terminal region, and ORF10 (Kotaki et al., 2021). This replicon was prepared based on the multiple ligation reactions *in vitro* of eight PCR amplicons; therefore, it was time-consuming and not useful. This replicon had other disadvantages in that it showed transient expression and its genome structure markedly differed from that of authentic SARS-CoV-2.

In the present study, we improved the usability of the previously reported replicon by cloning it into a highly stable vector, pCC2Fos (pCC2Fos-Δorf2-8 replicon). We also attempted to establish a stable replicon that is constantly sustained in cells. Although some groups reported the establishment of stable replicons, they were expressed in non-human-derived cells that affected the interferon response and they also lacked accessory or structural genes (Liu et al., 2022; Nguyen et al., 2021; Tanaka et al., 2022). These replicons may not necessarily reflect viral replication kinetics in the human body. Therefore, it is important to develop a more practical and stable replicon that has as many viral genes as possible and is sustained in interferon-competent, human-derived cells. Accordingly, we constructed new replicons, including other viral accessory and structural genes, and transfected them into Huh-7 cells with the expectation of imitating authentic viral replication under physiological conditions. We also selected replicon-transfected cells using puromycin for the establishment of stable replicons that are highly practical for the analysis of SARS-CoV-2 replication and drug screening in BSL-2 laboratories.

## 2. Materials and Methods

### 2.1. Virus and cell lines

A clinical SARS-CoV-2 isolate from Japan (JPN AI-I 004 strain; EPI_ISL_407084) was used to construct replicons. Chinese hamster ovary-K1 (CHO-K1) cells (ATCC: CCL-61) were maintained in Eagle’s minimal essential medium (EMEM) (NISSUI PHAMACEUTICAL CO., Tokyo, Japan) supplemented with 10% fetal bovine serum (FBS) (NICHIREI BIOSCIENCE INC., Tokyo, Japan) and 1% non-essential amino acid solution (NEAA) (NACALAI TESQUE, INC., Kyoto, Japan) at 37°C with 5% CO_2_. The human hepatoma cell line, Huh-7 was maintained in Dulbecco’s Modified Eagle Medium (DMEM) (FUJIFILM Wako Pure Chemical Corporation, Osaka, Japan) supplemented with 10% FBS and 1% NEAA.

### 2.2. Construction of the pCC2Fos replicon

To construct the pCC2Fos-Δorf2-8 replicon, Δorf2-8 replicon DNA was prepared according to a previously described method (Kotaki et al., 2021). Two complementary oligonucleotides including the T7 terminator and ZraI restriction site were annealed to prepare a 5’ side linker sequence (Supplementary Table 1). In the same manner, the 3’ side linker sequence including the Hepatitis D Virus ribozyme (HDVr) and ZraI site was prepared. These two linker sequences were connected to both termini of replicon DNA by ligation using 1,000 units of T4 DNA ligase [New England Biolabs (NEB), Ipswich, Massachusetts, USA] at 4°C overnight. Cloning into CopyControl™ pCC2Fos™ (Lucigen, Middleton, Wisconsin, USA) was conducted according to the manufacturer’s protocol using linker-connected replicon DNA as an insert.

To construct the pCC2Fos-Δorf2.4 and pCC2Fos-Δorf2 replicons, viral RNA extracted from the culture fluid of SARS-CoV-2-infected Vero E6 cells (provided by the National Institute of Infectious Disease, Japan) was reverse transcribed into cDNA by the SuperScript III First Strand Synthesis system (Thermo Fisher Scientific, Waltham, Massachusetts, USA) and cloned into the pCC2Fos-Δorf2-8 replicon by overlap PCR. TransforMaxTMEPI300TM Electrocompetent *Escherichia coli* (Lucigen) was transformed with these replicons using MicroPulser^™^ (Bio-Rad Laboratories, Inc., Hercules, California, USA), according to the manufacturer’s protocol.

To construct the pCC2Fos-puro replicon, the puromycin *N*-acetyltransferase (PAC) gene was amplified from pLenti CMV Puro DEST (w118-1) (Addgene #17452) by PCR, followed by the introduction of two silent mutations into the ZraI sites found in the PAC gene by inverse PCR. The PAC amplicon was inserted instead of S leaving its Transcription Regulatory Sequence (TRS).

To construct the RNA-dependent RNA polymerase (RdRp) mutant replicon, a fragment including the SARS-CoV-2 RdRp catalytic residue was subcloned into pUC19 (NEB) and the resulting pUC19-RdRp was mutated at the RdRp catalytic residue by inverse PCR using mutation-introducing primers (Supplementary Table 1). The intact RdRp of the pCC2Fos-Δorf2-8 replicon was replaced with mutated RdRp. All pCC2Fos replicons were extracted using the GenEluteTM HP Plasmid Miniprep Kit (Sigma Aldrich, St. Louis, Missouri, USA) and NucleoBond® Xtra Midi Plus (MACHEREY-NAGEL, Düren, Germany) for small and large scales, respectively.

### 2.3. In vitro *transcription*

The pCC2Fos replicon was digested with ZraI (NEB) and electrophoresed on 1% low melting point agarose (Nippon Gene Co., Tokyo, Japan). The bands corresponding to the replicons (23,304, 25,927, and 26,179 bp for the Δorf2-8, Δorf2.4, and Δorf2 replicons, respectively) were cut, cut gels were melted, and thermostable Δ-agarase was added (Nippon Gene). Replicon DNA in molten agarose was directly purified with phenol-chloroform-isoamyl alcohol (25:24:1), chloroform, and isopropyl alcohol precipitation. Precipitated replicon DNA was washed once with 70% ethanol, air-dried, and dissolved in nuclease-free water. Replicon RNA was then transcribed using the mMESSAGE mMACHINE T7 transcription Kit (Thermo Fisher Scientific) as previously described (Kotaki et al., 2021). After removing the DNA template following the manufacturer’s protocol, replicon RNA was extracted by TRIzol^™^ reagent (Thermo Fisher Scientific) and chloroform, followed by isopropyl alcohol precipitation. Precipitated RNA was washed once with 70% ethanol, dried by air, and dissolved in nuclease-free water.

### 2.4. Electroporation and luminescence quantification

Replicon RNA was electroporated using a NEPA21 electroporator (Nepagene, Chiba, Japan). Cells were trypsinized and washed once with phosphate-buffered saline (PBS) and twice with Opti-MEM (Thermo Fisher Scientific). In total, 1.0 × 10^6^ of washed cells were mixed with 5 μg of replicon RNA in 100 μl of Opti-MEM. Electric pulses were given by NEPA21. The parameters for CHO-K1 cells were as follows: voltage=145 V; pulse length=5 ms; pulse interval=50 ms; number of pulses=1; decay rate=10%; polarity+ as the poring pulse and voltage=20 V; pulse length=50 ms; pulse interval=50 ms; number of pulses=5; decay rate=40%; and polarity+/- as the transfer pulse. The parameters for Huh-7 cells were as follows: voltage=160 V; pulse length=5 ms; pulse interval= 50 ms; number of pulses=2; decay rate=10%; polarity+ as the poring pulse and voltage=20 V; pulse length=50 ms; pulse interval=50 ms; number of pulses=5; decay rate=40%; polarity +/- as the transfer pulse. After electroporation, cells were seeded at 1.5 × 10^4^ cells/120 µl/well on a 96-well plate. At various time points post-transfection, cells were lysed with 25 μl of the Nano-Glo HiBiT lytic detection system (Promega, Madison, USA) plus 25 μl of PBS. The luminescence signal was detected by CentroPRO LB962 (Berthold Technologies, Bad Wildbad, Germany). Seventy-two hours post-transfection (hpt), the medium was changed. Luminescence in the supernatant of Δorf2 replicon-transfected cells was detected using 50 µl of the supernatant and 50 µl of PBS-plus HiBiT lytic buffer solution described above.

### 2.5. Antiviral agent treatment

Huh-7 cells were electroporated with 5 µg of replicon RNA in the same manner as that described above and seeded at 1.5 × 10^4^ cells/120 µl/well on a 96-well plate. Immediately after electroporation, cells were treated with remdesivir at various concentrations or 0.2 % DMSO as the solvent control. Twenty-four hpt, the luminescence signal was measured as described above. The EC_50_ value was calculated using a four-parameter logistic regression model from GraphPad Prism 8 software (GraphPad Software Inc., Boston, MA)

### 2.6. Puromycin treatment

Huh-7 cells electroporated with 5 µg of puro replicon RNA were seeded at 1.5 × 10^4^ cells/120 µ/well on a 96-well plate. The HiBiT signal was measured at the indicated time points and the medium was changed at 72 hpt. To validate the effects of puromycin on the retention of replicons, puro replicon RNA-electroporated cells were seeded at 1.0 × 10^4^ cells/120 µl/well on a 96-well plate. At 24 hpt, the medium was changed to that containing various concentrations of puromycin or no puromycin, and the HiBiT signal was measured. The medium was changed again at 72 hpt.

### 2.7. Quantitative reverse transcription PCR (qRT-PCR)

After electroporation, transfected Huh-7 cells were seeded at 1.0 × 10^5^ cells/500 µl/well on a 24-well plate. Replicon RNA was extracted using NucleoSpin^®^RNA (MACHEREY-NAGEL) at the indicated time points and its copy number was measured using the THUNDERBIRD Probe One-step qRT-PCR Kit (TOYOBO CO., Osaka, Japan). CDC-approved primers (CDC_2019-nCoV_N2-F and CDC_2019-nCoV_N2-R) and a probe (CDC_2019-nCoV_N2-P) targeting the SARS-CoV-2 N gene were used. The probe contained a 6-carboxyfluorescein reporter dye at the 5′-end and a Black Hole Quencher (BHQ) at the 3′-end (Supplementary Table 1). An *in vitro* transcribed SARS-CoV-2 N gene was used as the RNA standard for qRT-PCR. The qRT-PCR reaction was set up in a 20-μl volume with 100 - 200 ng of RNA and was conducted using the CFX Connect Real-Time PCR System (Bio-Rad).

### 2.8. Immunofluorescent assay (IFA)

To detect membrane proteins and NSP8, Huh-7 cells were transfected with 5 μg of Δorf2-8, Δorf2.4, or Δorf2 replicon RNA and seeded on the Nunc™ Lab-Tek™ II Chamber Slide™ System (Thermo Fisher Scientific). At 48 hpt, transfected cells were fixed with 4% paraformaldehyde in PBS, permeabilized with 0.5% Triton-X, and blocked with 5% normal goat serum (Jackson ImmunoResearch Laboratories, Inc., West Grove, Pennsylvania, USA) in PBS. Rabbit anti-SARS-CoV-2 Membrane Glycoprotein polyclonal antibodies (ProSci, Poway, California, USA) and mouse anti-SARS-CoV-2 NSP8 mAb (5A10, GeneTex, Irvine, California, USA) were used in a single mixture for the primary antibody reaction. Goat anti-mouse IgG H&L conjugated with Alexa Fluor 568 (Abcam, Cambridge, UK) and goat anti-rabbit IgG H&L conjugated with Alexa Fluor 488 (Abcam) were used in a single mixture for the secondary antibody reaction. Cells were mounted with a mounting medium containing 4′,6-diamidino-2-phenylindole (DAPI) (Vector Laboratories, Burlingame, California, USA), which stained nuclei. To detect HiBiT, Huh-7 cells were transfected with 5 μg of Δorf2.4-puro replicon RNA and seeded on the chamber slide, as described above. At 24 hpt, transfected cells were fixed with 4% paraformaldehyde in PBS, permeabilized with 0.5% Triton-X, and blocked with 3% normal goat serum (Jackson ImmunoResearch Laboratories, Inc) in PBS. Mouse anti-HiBiT tag mAb (30E5, Promega) and goat anti-mouse IgG H&L conjugated with Alexa Fluor 568 (Abcam) were used for primary and secondary antibody reactions, respectively. Cells were mounted with a mounting medium containing DAPI (Vector Laboratories), which stained nuclei. Fluorescence images were acquired using the ZOE Fluorescent Cell Imager (Bio-Rad).

### 2.9. Statistical analysis

All statistical analyses were performed using GraphPad Prism 8 (GraphPad Software Inc.) with a two-way analysis of variance (ANOVA).

## 3. Results

### 3.1. Construction and characterization of the pCC2Fos-Δorf2-8 replicon

After the construction of the Δorf2-8 replicon according to the previously reported method (Kotaki et al., 2021), two linker sequences were connected to both ends of the replicon; one included the T7 terminator (T7-term) and the ZraI site, while the other included HDVr and the ZraI site. Cloning into pCC2Fos was performed using this modified replicon as an insert, and the resulting pCC2Fos-Δorf2-8 replicon was extracted (Figs. 1A and 1B). After digestion with ZraI, the Δorf2-8 replicon was purified and transcribed into replicon RNA. To verify replication, CHO-K1 cells were electroporated with replicon RNA and the HiBiT signal was measured at various time points, similar to the previous study (Kotaki et al., 2021). HiBiT kinetics were similar to that previously reported, with a peak at 48 hpt (Fig. 1C).

**Fig 1.**
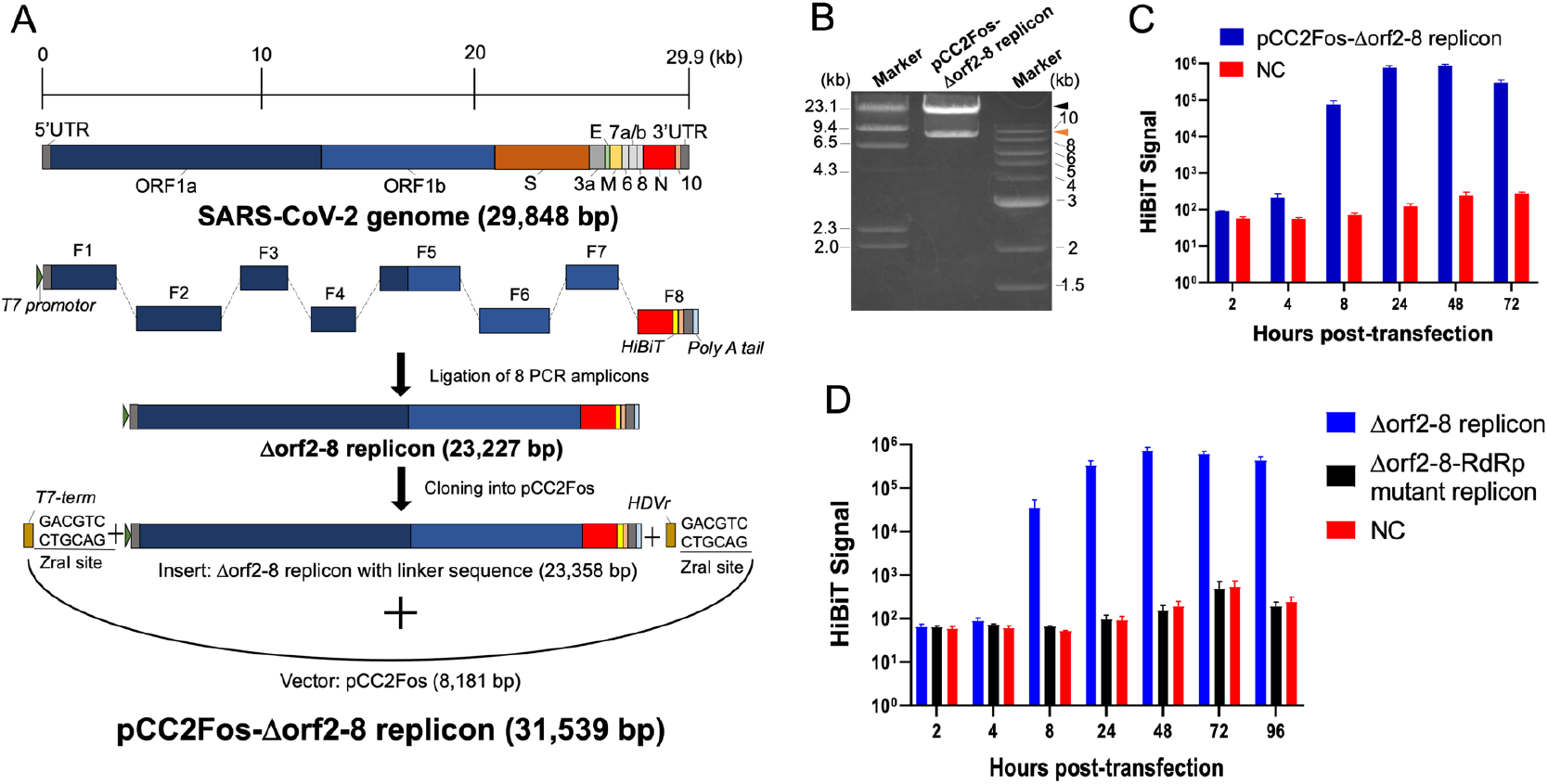
Construction and characterization of the pCC2Fos-Δorf2-8 replicon. (A) Schematic illustration of the pCC2Fos-based replicon. The Δorf2-8 replicon with linker sequences was cloned into pCC2Fos. (B) Electrophoretic image of the pCC2Fos-Δorf2-8 replicon after digestion with ZraI. The larger band (shown by a black arrowhead) is similar to the Δorf2-8 replicon (23,304 bp), while the smaller band (shown by an orange arrowhead) is similar to pCC2Fos (8,235 bp). (C) HiBiT kinetics of the pCC2Fos-Δorf2-8 replicon. CHO-K1 cells were electroporated with Δorf2-8 replicon RNA and the HiBiT signal was measured at the indicated time points. (D) HiBiT kinetics of the Δorf2-8 replicon and Δorf2-8-RdRp mutant replicon in Huh-7 cells. Huh-7 cells were electroporated with the Δorf2-8 replicon or Δorf2-8-RdRp mutant replicon, which theoretically does not replicate, and the HiBiT signal was measured at the indicated time points. For panels C and D, the averages and standard errors of at least 2 independent experiments with triplicate samples are shown.

### 3.2. Replication of replicons in the human hepatoma cell line, Huh-7

To compare the replication level of RNA replicons between a hamster-derived cell line (CHO-K1 cells) and human-derived cell line, Huh-7 cells, a human hepatic carcinoma cell line, were electroporated with the Δorf2-8 replicon. In Huh-7 cells, the HiBiT signal of the Δorf2-8 replicon peaked at 48 hpt, which was the same as that in CHO-K1 cells. Huh-7 cells were simultaneously electroporated with an RdRp-mutated replicon, which had two amino acid alterations in its catalytic residues: D760 and D761 were altered to A760 and A761 (Gao et al., 2020). The HiBiT signal of cells transfected with the Δorf2-8-RdRp mutant replicon was the same as that of the negative control because RdRp is a pivotal protein in viral replication (Fig. 1D). This result validated the RdRp-based replication of the Δorf2-8 replicon. Although the HiBiT peak time was similar between the two cell lines (48 hpt), a slightly sustained HiBiT signal was observed in Huh-7 cells at 72 and 96 hpt. At this point, replicons appeared to be easier to sustain in Huh-7 cells than in CHO-K1 cells. Therefore, we proceeded to the next experiments with Huh-7 cells as transfected cells.

### 3.3. Design of new replicons with other viral genes

We constructed two new replicons that possessed viral genes other than ORF1 and ORF9, and named them the Δorf2 replicon and Δorf2.4 replicon. The Δorf2 replicon is the structurally maximal replicon that possesses all viral genes, except for S, and is expected to simulate authentic viral replication (Fig. 2A). The Δorf2.4 replicon is the second most structurally maximal replicon that is lacking a structural gene found in the Δorf2 replicon (Fig. 2A). The M gene was maintained because it was previously reported to be associated with the anti-interferon response (Lee et al., 2022). These new replicons were constructed by the ligation of the pCC2Fos-Δorf2-8 replicon and other additional genes. The resulting pCC2Fos-Δorf2.4 replicon and pCC2Fos-Δorf2 replicon were confirmed by electrophoresis on an agarose gel (Fig. 2B) and a sequence analysis (data not shown). The difference between each band size on an agarose gel was not clear due to the limit of electrophoresis to separate bands on an agarose gel (Fig. 2B). Following the electroporation of these replicon RNAs into Huh-7 cells, their replication was confirmed by HiBiT luminescence (Fig. 2C) and their viral protein expression by IFA staining of non-structural protein 8 (NSP8) and a membrane protein that was not included in the Δorf2-8 replicon (Fig. 2D). The Δorf2.4 replicon showed similar HiBiT signal kinetics to the Δorf2-8 replicon, reaching its peak at 48 hpt; however, its intensity was approximately 2.3-fold lower than that of the Δorf2-8 replicon (Fig. 2C). The Δorf2 replicon exhibited different signal kinetics from the other replicons, showing its peak at 24 hpt (Fig. 2C). Since the HiBiT signal was detected in the supernatant of Δorf2 replicon-transfected cells, the production of virus-like particles (VLP) was suggested (Fig. 2E). These results indicated that the Δorf2 replicon was not promising due to poor replicon retention in cells for a stable replicon system. Regarding the Δorf2.4 replicon, although it showed a lower signal intensity, it still had the advantage of possessing other accessory genes and structural genes. Therefore, we selected the Δorf2-8 replicon and Δorf2.4 replicon as promising replicons for developing a stable replicon system.

**Fig. 2.**
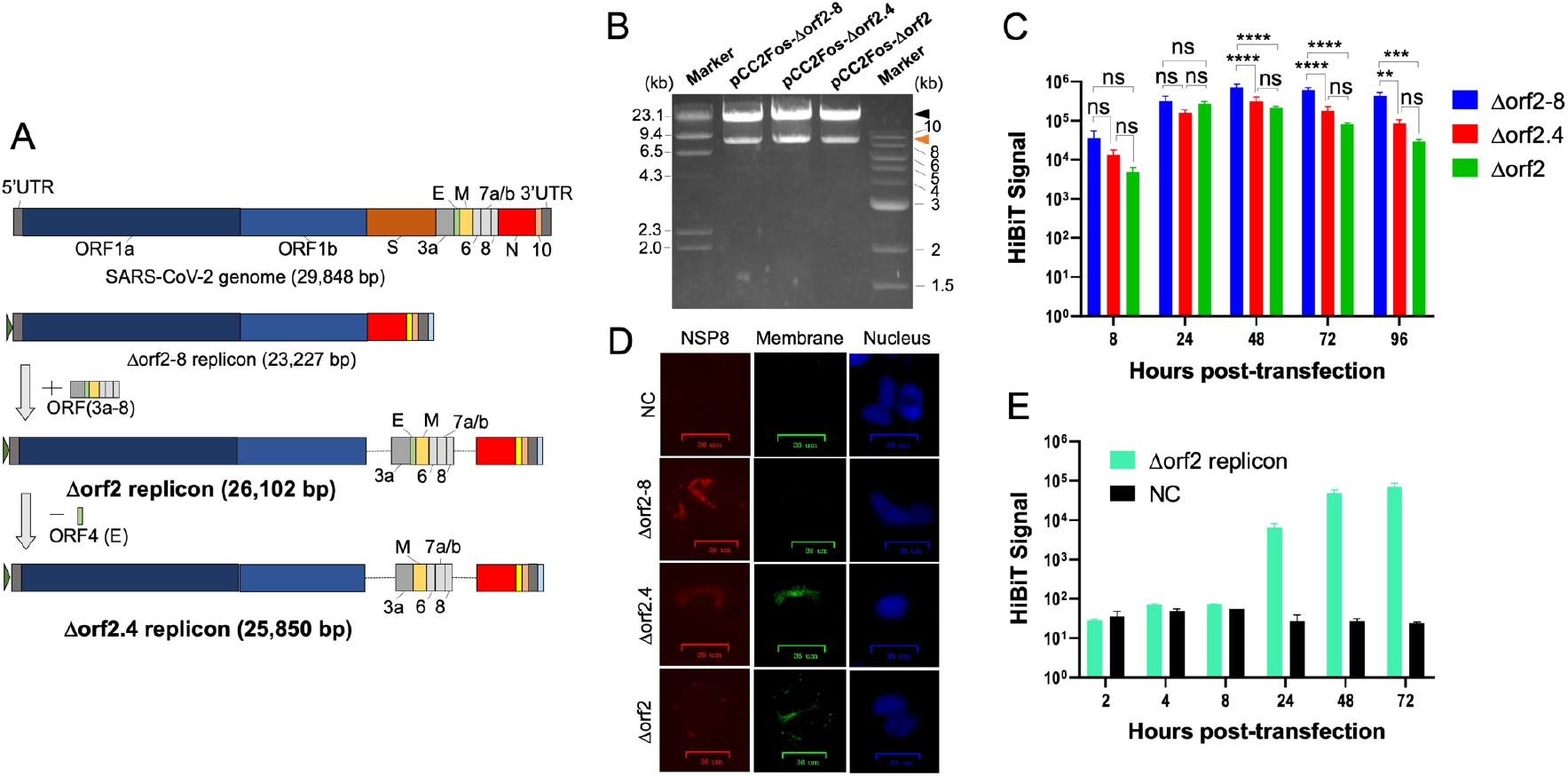
Construction and characterization of the Δorf2.4 replicon and Δorf2 replicon. (A) Schema of the two new replicons. The Δorf2 replicon lacks only S (the brown box), while the Δorf2.4 replicon lacks S and E (the green box). (B) Electrophoretic image of the pCC2Fos-Δorf2.4 replicon and pCC2Fos-Δorf2 replicon digested with ZraI. In each of the migrating images, the larger bands are similar to each replicon and the smaller bands to pCC2Fos. The theoretical band sizes of the replicons are 25,927 and 26,179bp for the Δorf2.4 replicon and Δorf2 replicon, respectively. (C) HiBiT signal kinetics of all three replicons. Huh-7 cells were electroporated with each replicon and the HiBiT signal was measured at the indicated time points. The significance of differences was assessed by a two-way ANOVA, considering p<0.05 as significant (**=p<0.01, ***=p<0.001, ****=p<0.0001). ns means Not Significant. (D) Immunofluorescent assay for viral protein detection. At 48 hpt, NSP8 was stained using the mouse anti-SARS-CoV-2 NSP8 antibody and goat anti-mouse IgG antibody conjugated with Alexa Fluor568 (red-colored cells), and the membrane was stained using the rabbit anti-SARS-CoV-2 membrane antibody and goat anti-rabbit IgG antibody conjugated with the Alexa Fluor 488 antibody (green-colored cells). The nucleus was stained by DAPI. (E) HiBiT luminescence in the supernatant of Δorf2 replicon-transfected cells. Huh-7 cells were electroporated with the Δorf2 replicon and HiBiT luminescence in its supernatant was measured at the indicated timepoints. For panels C and E, the averages and standard errors of at least 2 independent experiments with triplicate samples are shown.

### 3.4. Functional validation of replicons for antiviral agent screening

We assessed the applicability of two replicons, the Δorf2-8 and Δorf2.4 replicons, to antiviral agent screening. Immediately after the electroporation of the Δorf2-8 or Δorf2.4 replicon, Huh-7 cells were immediately treated with various concentrations of remdesivir, a representative SARS-CoV-2 replication inhibitor that is used in clinical settings (Beigel et al., 2020; Wang et al., 2020). At 24 hpt, the HiBiT signal of remdesivir-treated cells was measured. The HiBiT signal decreased in a dose-dependent manner, and EC_50_ values were calculated as 0.025 and 0.019 μM in cells transfected with the Δorf2-8 replicon and Δorf2.4 replicon, respectively (Figs. 3A and 3B). In Huh-7.5 cells, Huh-7-derived cells that are defective in interferon production, the EC_50_ value of remdesivir was 0.03 μM using an authentic virus (Ramirez et al., 2021). On the other hand, the EC_50_ value of remdesivir in Huh-7.5 cells was 0.0189 μM using a replicon (He et al., 2021). Collectively, these findings and the present results showed that the replication of replicons was inhibited by remdesivir at the same level as that in an authentic virus, with similar EC_50_ values. Therefore, the functionality of the Δorf2-8 and Δorf2.4 replicons for antiviral drug screening was validated.

**Fig. 3.**
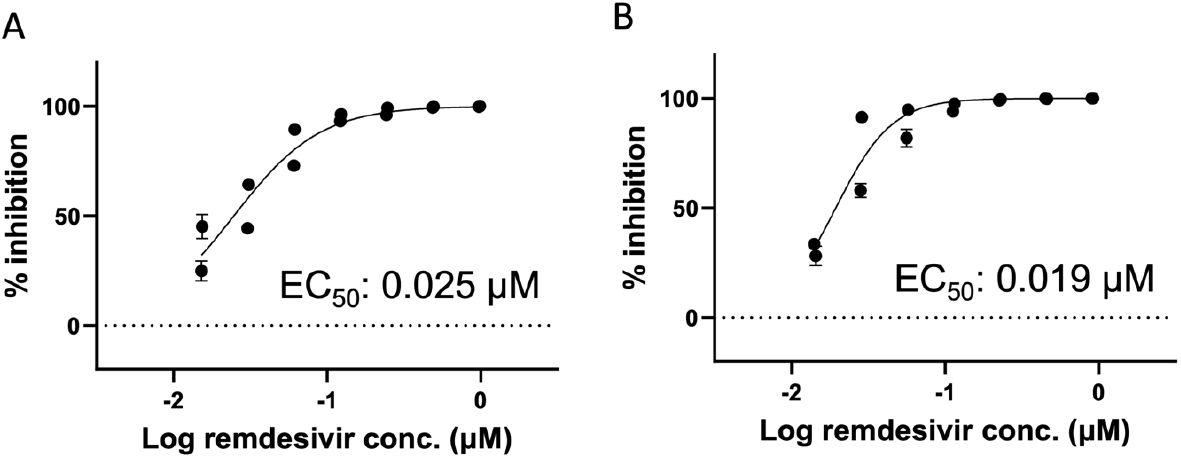
Regression curve of HiBiT luminescence by the remdesivir treatment. Immediately after the electroporation of Huh-7 cells with the Δorf2-8 replicon (A) or Δorf2.4 replicon (B), cells were treated with various concentrations of remdesivir. At 24 hpt, HiBiT luminescence was measured and a regression curve was created. EC_50_ values were calculated by GraphPad software. The averages and standard errors of 2 independent experiments with triplicate samples are shown.

### 3.5. Puromycin treatment of replicon-transfected cells to maintain the replication of replicons

To develop replicons that are sustained for a long time, we constructed a new replicon with a puromycin-resistant gene (puro replicon) and then subjected cells transfected with puro replicons to a treatment with puromycin. The PAC gene was inserted at the ORF2 cassette of two replicons, the Δorf2-8 and Δorf2.4 replicons. The resulting Δorf2-8-puro and Δorf2.4-puro replicons were electroporated into Huh-7 cells to confirm replicon replication after PAC gene incorporation (Fig. 4A). The HiBiT signal of the Δorf2.4-puro replicon was similar to that of the Δorf2.4 replicon. However, the HiBiT signal of the Δorf2-8-puro replicon was approximately 50-fold lower than that of the Δorf2-8 replicon, and was significantly weaker than that of the Δorf2.4-puro replicon (Fig. 4A). Furthermore, a comparison of replicon RNA copy numbers between the two puro replicons revealed a significant difference at the RNA level, indicating a difference in replication capacity (Fig. 4B). Therefore, we selected the Δorf2.4-puro replicon as a promising replicon that is sustained in cells for a long time. We then treated Δorf2.4-puro replicon-transfected Huh-7 cells with puromycin. At 24 hpt, the HiBiT signal was measured to set a baseline and the medium was changed to new medium containing various concentrations of puromycin (1.0, 0.4, 0.16, 0.08, and 0 μg/mL) (Fig. 4C). The HiBiT signal of puromycin-treated cells was measured at the indicated time points. A significantly stronger signal was detected at 72 hpt and thereafter in cells treated with 1.0 µg/mL of puromycin than in non-treated cells. Similar results were obtained for cells treated with 0.4 μg/mL of puromycin, but without a significant difference at any time point. Regarding cells treated with 0.16 or 0.08 μg/mL of puromycin, no significant differences were observed from cells not treated with puromycin (data not shown). We also attempted to clone cells stably expressing replicons using 0.4 and 1.0 μg/mL of puromycin, but have not yet succeeded. We then conducted IFA to examine the transfection efficiency of the Δorf2.4-puro replicon. The expression of the HiBiT protein was detected using anti-HiBiT antibodies (Fig. 4D). The number of HiBiT-positive cells (red-colored cells) was counted and transfection efficiency was calculated as 2.7%. This score was close, but lower than that reported in Huh-7.5 cells (3-4 %) using another larger replicon (He et al., 2021).

**Fig. 4.**
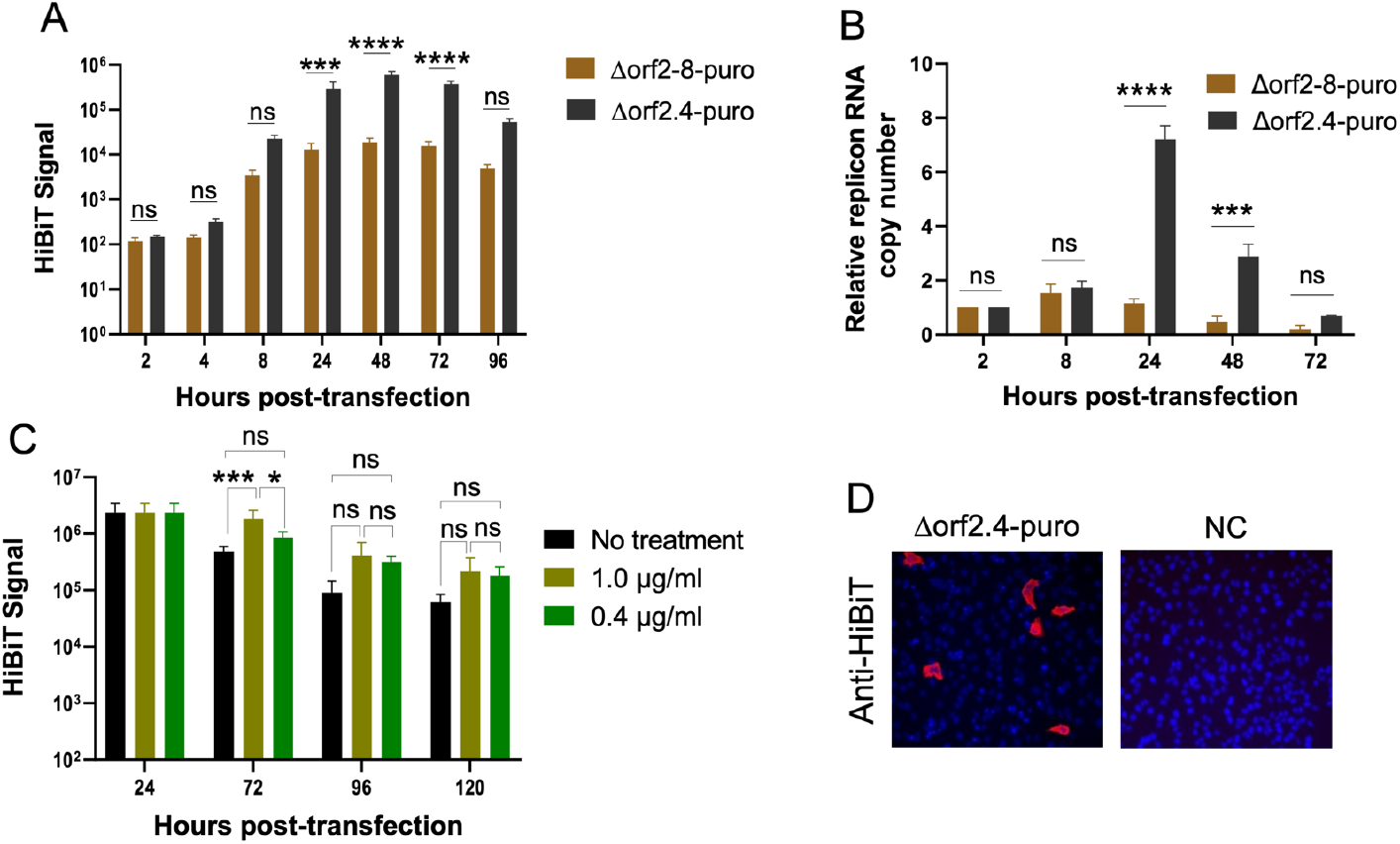
Puromycin treatment. (A) HiBiT kinetics of two puro replicons without puromycin pressure. Huh-7 cells were electroporated with the Δorf2-8 puro replicon or Δorf2.4-puro replicon, and the HiBiT signal was measured at the indicated timepoints. (B) Kinetics of replicon RNA copy numbers. The total RNA of Huh-7 cells electroporated with the puro replicon was extracted and the expression of the N gene was quantified by qRT-PCR. The graph shows the relative replicon (N) RNA copy number normalized by the copy number at 2 hpt. (C) HiBiT luminescence under the puromycin treatment. Huh-7 cells were electroporated by the Δorf2.4-puro replicon and treated with various concentrations of puromycin at 24 hpt. The medium was changed again at 72 hpt. The signal at 24 hpt is the same between samples considering the baseline. (D) Immunofluorescent assay (IFA) for the detection of HiBiT. The expression of HiBiT was detected with the anti-HiBiT tag mAb and goat anti-mouse IgG antibody conjugated with Alexa Fluor 568 (red-colored cells). The nucleus was stained by DAPI. For panels A, B and C, the averages and standard errors of at least 2 independent experiments with triplicate samples are shown. The significance of differences was assessed by a two-way ANOVA considering p<0.05 as significant (**=p<0.01, ***=p<0.001, ****=p<0.0001). ns means Not Significant.

## 4. Discussion

We herein constructed a series of SARS-CoV-2 RNA replicons using the highly stable vector, pCC2Fos. To construct a template plasmid of the SARS-CoV-2 replicon, other research groups frequently used a BAC vector, a well-known stable vector (He et al., 2021; Zhang et al., 2021a; Zhang et al., 2021b). The present study is the first to report a pCC2Fos-based replicon and the results obtained provide novel insights into cloning strategies for replicons including emerging viruses in the future. This pCC2Fos replicon is more convenient than the previously reported PCR amplicon-based replicon system in terms of preparation times and the risk of unexpected mutations. It also enabled us to add other virus genes, exogenous genes, and mutations of interest into replicons, which will lead to the development of more practical replicon systems and efficient mutation analyses. We herein constructed two new replicons, the Δorf2 and Δorf2.4 replicons in addition that that previously reported (Kotaki et al., 2021) by adding other viral genes, with the expectation of closely replicating the mechanism of the authentic virus and increasing the sustainability of RNA replication. Although the replication of these replicons was confirmed, as observed in HiBiT signal kinetics (Fig. 3C), these modifications did not improve replicon sustainability. The reduced intensity of the two new replicons was attributed to the template lengths in RNA transcription and replication. Since the addition of other viral genes increased the template length by approximately 3 kb (2623 bp for the Δorf2.4 replicon and 2875 bp for the Δorf2 replicon), the efficiency of *in vitro* transcription and minus-sense RNA genome replication were lower than those by the Δorf2-8 replicon. Since our replicons are equipped with a HiBiT reporter gene fused to the C terminus of N, an increase in not fully transcribed replicon RNA leads to a signal reduction. Another possibility is that the existence of other ORFs between ORF1 and N affects the efficiency of subgenomic RNA (sgRNA) transcription. Transcripts from the Δorf2-8 replicon only consisted of two RNAs, N-sgRNA and genomic replicon RNA. On the other hand, more sgRNAs, other than S-sgRNA and E-sgRNA, were transcribed from the Δorf2.4 replicon, while all sgRNAs other than S-sgRNA were transcribed from the Δorf2 replicon. This may result in the poorer recruitment of the replication complex to the TRS of ORF1, namely, the reduced replication of replicons, than that of the Δorf2-8 replicon. Other factors may be attributed to the effects of accessory proteins. The ORF3a protein included in the Δorf2 and Δorf2.4 replicons has been reported to induce apoptosis via the caspase-3 and -8 pathways, which may cause cell death and a reduced HiBiT signal (Ren et al., 2020). Nevertheless, our Δorf2.4 replicon is still advantageous with functionality for drug screening. Our Δorf2 replicon, which has not yet been functionally validated, also has the advantage of being able to produce VLP. It enabled us to measure HiBiT luminescence without cell lysis. Other groups reported that replicons lacking only the S gene are applicable to the production of single-cycle infectious particles by the trans-complementation of the SARS-CoV-2 spike or other viral surface glycoproteins, such as the vesicular stomatitis virus glycoprotein (Cheung et al., 2022; Malicoat et al., 2022). The further application of our Δorf2 replicon to the production of single-cycle virus is of interest.

After the incorporation of the PAC gene into replicons, the Δorf2-8 puro replicon showed a reduced signal. Since no mutations were found by a sequencing analysis of the whole replicon genome (data not shown), a possible reason for this phenomenon is the effect of an additional ORF cassette. As described above, an increase in the number of ORF may weaken the replication of replicons. However, this phenomenon was not applicable to the Δorf2.4-puro replicon. We speculate that some accessory proteins confer beneficial effects on RNA replication, enhancing the recruitment of the polymerase complex to the ORF1 TRS sequence. A previous study reported that the deletion of ORF7a and ORF8 resulted in low viral titers of a recombinant live virus (Silvas et al., 2021). These accessory proteins are well-known interferon antagonists and reduced viral titers may be attributed to a defect in the anti-interferon response (Lei et al., 2020; Xia et al., 2020). However, there may be another effect on replication that has not yet been identified. The replication of the Δorf2.4-puro replicon was validated and enhanced by puromycin. However, we have not successfully established a stable Δorf2.4-puro replicon.

To the best of our knowledge, only three studies have reported the establishment of cells that stably harbor replicons (Liu et al., 2022; Nguyen et al., 2021; Tanaka et al., 2022). Two of them are DNA-based replicons and the other is an RNA-based replicon. A number of factors may be contributing to replicon stability. Regarding the structures of these three replicons, the two DNA-based replicons consisted of the same viral genes as our Δorf2-8 replicon and the RNA-based replicon had a similar structure to our Δorf2.4 replicon, consisting of all viral genes, except for S, E, and M. On the other hand, these replicons all possessed a neomycin-resistant gene as a selection marker, which is a distinct structural difference from our replicons. Since the selection pressure of puromycin is stronger than that of neomycin, puromycin is superior for short-term selection. However, this may work negatively in replicon-transfected cells, among which rare cells possessed puromycin resistance, as shown in Fig. 4D, and the low selection pressure of neomycin may result in a more stable replicon (Liu et al., 2022; Nguyen et al., 2021; Tanaka et al., 2022). Although selection markers are common among stable replicons, its inserted position differs. Since sgRNA transcription levels differ in every ORF and cells to be transfected, it is important to select an ORF cassette that is the most abundantly transcribed in cells for replacement with a selection marker (Jin et al., 2021). The physiological characteristics of transfected cells to be transfected are also critical. Stable replicons were established in interferon-deficient cells (BHK-21 cells and VeroE6 cells); therefore, replicons may be easily sustained under conditions without an interferon response (Liu et al., 2022; Nguyen et al., 2021; Tanaka et al., 2022). However, replicons are not necessarily stable in interferon-deficient cells. A previous study reported that a stable replicon in BHK-21 cells was not stable in Huh-7.5 cells, which lacked a RIG-I receptor and interferon response (Liu et al., 2022). Therefore, interferon signaling does not appear to be the only factor affecting replicon stability; however, the reason responsible remains unknown. In addition to the deficiency of interferon, the reported stable replicons are present in non-human cells. Therefore, the establishment of stable interferon-competent replicons in human-derived cells is more beneficial because they reflect viral kinetics in the human body.

The present results show that the insertion of the PAC gene instead of the S gene in Huh-7 cells is not optimal for stable replicons. Since the sgRNA of the N gene is the most abundant in Huh-7 cells, an additional selection marker to N may lead to better outcomes (Jin et al., 2021). However, the Δorf2.4-puro replicon has benefits over other stable replicons in that it replicates more similarly to authentic SARS-CoV-2 than DNA-based replicons, which cause DNA to RNA transcription in transfected cells that does not occur in SARS-CoV-2 infection. It also possesses more viral genes with validated functionality for antiviral drug screening. We also observed the sustained replication of the replicon under puromycin pressure. The present results provide important insights for the development of more practical and stable replicons that replicate by a closer mechanism to authentic SARS-CoV-2 than previously reported replicons. They will contribute to further research on SARS-CoV-2 replication and the more efficient discovery of SARS-CoV-2 replication inhibitors in BSL-2 facilities.

## CRediT authorship contribution statement

Shunta Takazawa: Methodology, Formal analysis, Investigation, Writing - Original Draft. Tomohiro Kotaki: Conceptualization, Methodology, Formal analysis, Writing - Review & Editing, Supervision. Satsuki Nakamura: Investigation, Writing - Review & Editing. Chie Utsubo: Resources, Writing - Review & Editing, Funding acquisition. Masanori Kameoka: Conceptualization, Resources, Writing - Review & Editing, Supervision, Project administration, Funding acquisition.

## Acknowledgments

We thank Dr. Masayuki Saijo of the National Institute of Infectious Diseases for providing RNA of the SARS-CoV-2 JPN AI/I 004 strain. The manuscript was proofread by Medical English Service, Kyoto, Japan.

## Supplementary materials

Supplementary Table 1. Sequences of primers and the probe.

