## Supplemental Table 1 for "Construction of Fosmid-based SARS-CoV-2 replicons for antiviral drug screening and replication analyses in biosafety level 2 facilities"

**Supplementary Table 1.** Sequences of primers and the probe.

| Name | Sequence (5’-3’) | Annotation |
| --- | --- | --- |
| HDVr-ZraI For | AAAAGGGTCGGCATGGCATCTCCACCTCCTCGCGGTCCGACCTGGGCTACTTCGGTAGGCTAAGGGAGAAGGACGTC | Annealed primers for the preparation of the 3’ side linker  sequence |
| HDVr-ZraI Rev | GACGTCCTTCTCCCTTAGCCTACCGAAGTAGCCCAGGTCGGACCGCGAGGAGGTGGAGATGCCATGCCGACCC |  |
| T7-term-ZraI For | CTAGCTATTCCCCTTGGGGACTCTAAACGGGTCTTGAGGGGTTTTTTGGACGTC | Annealed primers for the preparation of the 5’ side linker  sequence |
| T7-term-ZraI Rev | ATCAGACGTCCAAAAAACCCCTCAAGACCCGTTTAGAGGCCCCAAGGGGTTATGCTAG |  |
| RdRp mutation For | CTCTCTGCCGCTGCTGTTGTGTGTTTC | Primers for the RdRp mutation. |
| RdRp mutation Rev | AACAGCAGCGGCAGAGAGTATCATCA |  |
| CDC_2019-nCoV_N2-F | TTACAAACATTGGCCGCAAA | Forward primer for N detection |
| CDC_2019-nCoV_N2-R | GCGCGACATTCCGAAGAA | Reverse primer for N detection |
| CDC_2019-nCoV_N2-P | [FAM]ACAATTTGCCCCCAGCGCTTCAG[BHQ] | Probe for N detection |
| Puro ZraI mut1 Fwd | GCGACGATGTGCCCAGGGCCGTACGC | Primers for the mutation of two ZraI sites in *Puromycin N-acetyltransferase* |
| Puro ZraI mut1 Rev | CCCTGGGCACATCGTCGCGGGTGGCG |  |
| Puro ZraI mut2 Fwd | CCGCCGATGTGGAGGTGCCCGAAGGAC |  |
| Puro ZraI mut2 Rev | GCACCTCCACATCGGCGGTGACGGTG |  |
| SARS-CoV-2 ORF1b For | CGGTATAAATTAGAAGGCTATGCCTTCGAACA | Forward primer including the BstBI site |
| SARS-CoV-2 ORF1b Rev | TTAGTTGTTAACAAGAACATCACTAG |  |
| SARS-CoV-2 ORF3 For | ATGTTCTTGTTAACAACTAAACGAACTTATGGATTTG |  |
| SARS-CoV-2 ORF3 Rev | GAAAAACTAATATAATATTTAGTTCGTTTACAAAGGCACGCTAGTAGTCGTC |  |
| SARS-CoV-2 M For | ACGAACTAAATATTATATTAGTTTTTCTGTTTGG |  |
| SARS-CoV-2 N Rev | TTAATTGGAACGCCTTGTCCTCGAGG | Reverse primer including the XhoI site |
| SARS-CoV-2 1b-Puro | ATGTTATTGTTAACAACTAAACGAACAATGACCGAGTACAATCC | Forward primer for *PAC* amplification |
| SARS-CoV-2 Puro-3a | CAAATCCATAAGTTCGTTTAGGCACCGGGC | Reverse primer for *PAC* amplification |
| SARS-CoV-2 ORF3-F for Puro | CTAAACGAACTTATGGATTTGTTTATGAG | Forward primer to insert *PAC* into the Δorf2.4 replicon |
| SARS-CoV-2 Puro-N | TTCGTTTAGGCACCGGGCTTGC | Reverse primer for *PAC* amplification |
| SARS-CoV-2 N-F for Puro | CCGGTGCCTAAACGAACAAACTAAAATGTC | Forward primer to insert *PAC* into the Δorf2-8 replicon |
